# Accelerating Cancer Vaccine Development for Human T-Lymphotropic Virus (HTLV) Using a High-Throughput Molecular Dynamics Approach

**DOI:** 10.1101/2023.06.07.544070

**Authors:** Abu Tayab Moin, Nurul Amin Rani, Md. Asad Ullah, Rajesh B. Patil, Tanjin Barketullah Robin, Nafisa Nawal, Talha Zubair, Syed Iftakhar Mahamud, Mohammad Najmul Sakib, Nafisa Nawal Islam, Md. Abdul Khaleque, Nurul Absar, Abdullah Mohammad Shohael

## Abstract

Human T-lymphotropic virus (HTLV), a retrovirus belonging to the oncovirus family, has long been linked to be associated with various inflammatory and immunosuppressive disorders. To combat the devastating impact of this virus, our study employed a reverse vaccinology approach to design a multi-epitope-based vaccine targeting the highly virulent subtypes of HTLV. We conducted a comprehensive analysis of the molecular interactions between the vaccine and Toll-like receptors (TLRs), providing valuable insights for future research on preventing and managing HTLV-related diseases and any possible outbreaks. The vaccine was designed by focusing on the envelope glycoprotein gp62, a crucial protein involved in the infectious process and immune mechanisms of HTLV inside the human body. Epitope mapping identified T cell and B cell epitopes with low binding energies, ensuring their immunogenicity and safety. Linkers and adjuvants were incorporated to enhance the vaccine’s stability, antigenicity, and immunogenicity. Two vaccine constructs were developed, both exhibiting high antigenicity and conferring safety. Vaccine construct 2 demonstrated expected solubility and structural stability after disulfide engineering. Molecular docking analyses revealed strong binding affinity between the vaccine construct 2 and both TLR2 and TLR4. Molecular dynamics simulations indicated that the TLR2-vaccine complex displayed enhanced stability, compactness, and consistent hydrogen bond formation, suggesting a favorable affinity. Contact analysis, Gibbs free energy landscapes, and DCC analysis further supported the stability of the TLR2-vaccine complex, while DSSP analysis confirmed stable secondary structures. MM-PBSA analysis revealed a more favorable binding affinity of the TLR4-vaccine complex, primarily due to lower electrostatic energy. In conclusion, our study successfully designed a multi-epitope-based vaccine targeting HTLV subtypes and provided valuable insights into the molecular interactions between the vaccine and TLRs. These findings should contribute to the development of effective preventive and treatment approaches against HTLV-related diseases.

## 1. Introduction

Human T-lymphotropic viruses (HTLV) belong to the human retroviruses family, specifically the genus Deltaretrovirus, and they infect the human body, leading to various diseases such as adult T-cell leukemia (ATL). HTLV-1, HTLV-2, HTLV-3, and HTLV-4 are the four major subtypes of HTLV that have been identified so far. These group of viruses not only cause malignancy but also other inflammatory and immunosuppressive diseases (1, 2). HTLVs are enveloped viruses with a diameter ranging from 80 to 100 nm and within the HTLV virions, the reverse transcriptase (RT), integrase, protease viral enzymes, and capsid proteins form complexes with the two covalently bonded genomic RNA strands (3, 4). The proviral genome of HTLVs is approximately 9kb in length and is flanked by 5’ and 3’ long terminal repeats (LTRs), which consist of U3, R, and U5 regions. These regions facilitate viral integration into the host genome and contain promoter elements with regulatory sequences for viral transcription (5, 6).

Among the members of HTLVs, the pathogenic roles of HTLV-1 and HTLV-2 in the human body are well-studied to date compared to HTLV-3 and HTLV-4, and these group of viruses are transmitted through bodily fluids like semen, blood and milk (2). HTLV-1, in particular, is the most life-threatening member of this viral family, causing fatal ATL in approximately 5% of the infected cases that characterizes high blood circulation, lymph node swelling and immunosuppression (7, 8). Of the 4 types of ATL, acute ATL is most aggressive form of leukemia that results in median survival of patients less than one year. Moreover, this specific strain can also cause HTLV-1-associated myelopathy/tropical spastic paraparesis (HAM/TSP) that results in the progressive weakness of both legs and 50% of untreated patients are expected to lose their ability to walk within 10 years of initial diagnosis. HTLV-2 was first detected in a patient with hairy cell leukaemia and is considered less pathogenic when compared to HTLV-1 (9). HTLV-2 has arguably been associated to the development of HAM/TSP in infected patients like HTLV1 (10). Moreover, HTLV-2 was also found to be associated with the onset of neuropathic disorders, including meningitis and chronic inflammatory demyelinating polyneuropathy (CIDP) and affecting patients clinical outcome negatively (11, 12). The global impact of HTLV-1 and HTLV-2 infections is alarming, with an estimated 5-10 million people infected worldwide (13). HTLV-1 is highly prevalent in indigenous populations in Japan and South America, while HTLV-2 is primarily found in Central and South America (14).

HTLV-3 and HTLV-4 have been reported in isolated forest areas in the Republic of Cameroon, however, their potential risk to the human health is still under investigation (15). That said, these two strains possess similar genomic composition and ancestral relationship with HTLV-1 and HTLV-2 (16). Additionally, studies found that subtle genomic reconstruction of HTLV-3 can mount the virus strain to produce virulent particles (17) suggesting that the possible pathogenic roles of HTLV-3 and 4 cannot be entirely neglected.

Despite the HTLV-related diseases possess higher threat to the global public health, these group of diseases remain highly underrepresented in terms of developing an effective countermeasure through robust scientific research. HTLV-associated malignancies are often treated with conventional chemotherapy, bone marrow transplantation or monoclonal antibodies but these approaches often result in poorer outcome (8). Additionally, although there is some antiretroviral therapy which has shown some efficacy in suppressing viral replication of HTLVs, currently, there is no licensed vaccine available for HTLVs that may aid in curing a mass infected individuals during any possible outbreaks or epidemics (18).

However, developing a vaccine against HTLVs has been challenging due to the ability of the virus to integrate its genetic material into the host genome, making it difficult to target with a vaccine. Additionally, the presence of a few distinct viral proteins that interfere with the immune system and the variability of the virus further complicate vaccine development (11, 19). Hence, developing a vaccine targeting the effective virulent protein against HTLVs is crucial due to their intricate mechanism of infection and evading the host defense system. Study suggests that candidates able to boost anti-HTLV-envelope-glycoprotein antibodies may have potential roles in combating HTLV infection (20).

This study focused on targeting the Envelope Glycoprotein GP62 of the viruses as a potential antigen for the subunit vaccine. Glycoproteins, located on the outer layer of the viral envelope, are easily recognized by the immune system. Enveloped glycoprotein which is a surface protein (SU) of HTLV attaches the virus to the host cell by binding to its receptor, while the transmembrane protein (TM) undergoes a conformational change upon this interaction with the host cell, activating its fusogenic potential and leading to membrane fusion at the host cell plasma membrane. This fusion process allows the viral nucleocapsid to enter the cytoplasm of the host cell (21, 22).

Finally, the aim of the study is to design an effective polyvalent subunit vaccine against the major causative agents of HTLV-related diseases i.e., HTLV-1, HTLV-2 using high-throughput bioinformatics strategies. Additionally, given that the pathogenic mechanism and virulence of other subtypes i.e., HTLV-3 remain under investigation, we also incorporated HTLV-3 subtype in our experimental design. However, since the virulent protein sequence of HTLV-4 is yet not reported in public repository, we didn’t consider HTLV-4 in our study. Despite the challenges in vaccine development, this research aims to provide molecular insights into how the designed subunit vaccine can counteract viral infections. The results obtained from this study will facilitate further in vitro and in vivo validations, advancing the understanding and potential implementation of the HTLV vaccines.

## 2. Methods

The stepwise methodology of the entire study is depicted in **Supplementary Figure S1** as a flowchart.

### 2.1. Strain and protein selection with biophysical property analyses

From NCBI (https://www.ncbi.nlm.nih.gov/) database, three distinct HTLV strains, namely, HTLV-1, HTLV-2, and HTLV-3, were identified, and a virulent protein named Envelope glycoprotein gp62 was chosen for further epitope mapping. The target protein sequences of the chosen strains were then retrieved in FASTA format from the UniProt database (https://www.uniprot.org). The chosen protein sequence was then uploaded to the online antigenicity tool VaxiJen v2.0 (http://vaxijen/VaxiJen/VaxiJen.html) to examine the antigenic property (23). Transmembrane topology was evaluated by using the TMHMM-2.0 server (https://services.healthtech.dtu.dk/service.php?TMHMM-2.0). By using the ExPASy ProtParam server (https://web.expasy.org/protparam//), we examined various physicochemical characteristics of the protein (24). To find the conservancy pattern, the homologous sequence sets of the selected antigenic proteins were obtained from the NCBI database using the BLASTp program.

### 2.2. Epitope mapping analyses and vaccine construction

We utilized the Immune Epitope Database (IEDB) online tool (https://www.iedb.org/) to predict T-cell and B-cell epitopes of selected protein sequences, while keeping the default parameters. The full reference set of human leukocyte antigen (HLA) alleles was selected to predict MHC class I-restricted CD8+ cytotoxic T-lymphocyte (CTL) epitopes using the NetMHCpan EL 4.0 prediction method suggested by IEDB (http://tools.iedb.org/mhci/) (25). For MHC class II-restricted CD4+ helper T-lymphocyte (HTL) epitopes, we used the IEDB-recommended 2.22 prediction method with the full reference set of HLA alleles (http://tools.iedb.org/mhcii/). To predict linear B-cell epitopes (LBL) of the chosen protein, we used the BepiPred technique 2.0 with the default parameters (http://tools.iedb.org/bcell/), and selected the highest-scoring LBL epitopes as prospective candidates for further analysis.

After the initial epitope prediction, we determined epitope antigenicity using the VaxiJen v2.0 server, predicted transmembrane topology using the TMHMM 2.0 server, and assessed allergenicity using the AllergenFP (https://ddg-pharmfac.net/AllergenFP/) and AllerTOP (https://www.ddg-pharmfac.net/AllerTOP/) servers. We also evaluated toxicity using the ToxinPred server (http://crdd.osdd.net/raghava/toxinpred/) (26). The HTL epitopes’ capacity to induce IFN-g, IL-4, and IL-10 was predicted using the IFNepitope (http://crdd.osdd.net/raghava/ifnepitope/), IL4pred (http://crdd.osdd.net/raghava/il4pred/), and IL10pred (http://crdd.osdd.net/raghava/IL-10pred/) servers, respectively (27–29). In addition, the epitopes’ conservancy was evaluated by generating a multiple sequence alignment using version 3.0 of the CLC Drug Discovery Workbench 3 software (30). Epitopes with the highest potential for vaccine construction, based on their high antigenicity, non-toxicity, non-allergenicity, and conservancy, were chosen. To create the vaccine, these epitopes were combined with adjuvants like PADRE and human beta-defensin (hBds), and selective linkers such as EAAAK, AAY, and GPGPG.

### 2.3. Analyses of the biophysical and structural properties of the vaccine

To ensure the safety and efficacy of the vaccine, antigenicity and allergenicity analyses were conducted. The solubility of the vaccine construct upon expression in Escherichia coli was evaluated using the Protein-Sol server (https://protein-sol.manchester.ac.uk/) (31). The biophysical characteristics of the vaccine constructions, including isoelectric pH, aliphatic and instability index, GRAVY values, hydropathicity, anticipated half-life, and other characteristics were assessed using the ProtParam tool of the ExPASy server. The secondary structure of the final multi-epitope vaccine was predicted using two servers, SOPMA (https://npsa-prabi.ibcp.fr/cgi-bin/npsa_automat.pl?page=/NPSA/npsa_sopma.html) and PSIPRED (http://bioinf.cs.ucl.ac.uk/psipred/) (32). The trRosetta server (https://yanglab.nankai.edu.cn/trRosetta/) was used to modulate the tertiary structure of the vaccine constructs (33, 34), which was later refined by the GalaxyRefine module of the GalaxyWEB server (http://galaxy.seoklab.org/) (35). The improved models were then validated through the PROCHECK server (https://saves.mbi.ucla.edu/) for Ramachandran plots and ERRAT score plots, and ProSA-web server (https://prosa.services.came.sbg.ac.at/prosa.php) for Z score plots (36–38).

### 2.4. Molecular docking analyses

Initially, a molecular docking analysis was conducted to investigate the binding affinity between the CTL epitopes and HLA alleles. For the docking study with T cell epitopes, HLA-A11:01 and HLA-DRB104:01 were selected. The PEP-FOLD Peptide Structure Prediction server was used to generate the 3D structures of the best T cell epitopes. The H-dock server (http://hdock.phys.hust.edu.cn/) was then utilized to perform the molecular docking analysis, demonstrating that the proposed epitopes could interact with at least one MHC molecule with minimum binding energy (39–41). Subsequently, another molecular docking analysis was carried out between the vaccine constructs and TLR2 and TLR4 receptors to predict their binding affinities and interaction patterns. The TLR2 and TLR4 receptor structures were obtained from the RCSB PDB database, and the refined 3D structure of the multi-epitope construct served as the ligand. The binding affinity between the vaccine construct and TLR2 and TLR4 was calculated using the ClusPro 2.0 server and H-dock server (https://cluspro.bu.edu/login.php) (42). The best-docked complex was identified based on the lowest energy-weighted score and docking efficiency.

### 2.5. Molecular dynamics simulation studies

The docked complexes of respective vaccine constructs with TLR2 and TLR4 were subjected to 100 ns molecular dynamics (MD) simulations with Gromacs 2020.4 (43) package using the resources of HPC cluster at Bioinformatics Resources and Applications Facility (BRAF), C-DAC, Pune. TLR2 and TLR4 have four chains each and the topologies for these chains and vaccine chains were prepared using the CHARMM-36 force field parameters (44, 45). The complexes of TLR2 and TLR4 with respective vaccine constructs were solvated with the TIP3P water model (46) after positioning them in a box of dodecahedron unit cells, keeping the boundary of system 1 nm from the edges of the box. The charges on each solvated system were neutralized where the TLR2 complex and the TLR4 complex required the addition of 8 and 16 sodium counter-ions, respectively. Further, the energy minimization was performed with the steepest descent algorithm employing the force constant threshold of 100 kJ mol^-1^ nm^-1^. The equilibration of each system for 1 ns was then carried out at constant volume and constant temperature (NVT) conditions where the temperature of 300 K was achieved with a modified Berendsen thermostat (47) and at constant volume and constant pressure (NPT) conditions where 1 atm pressure was achieved with Berendsen barostat (48). The production phase MD simulations of 100 ns were performed on each equilibrated system without any restraints on the chain of TLR2 and TLR4. During production phase, MD simulations the temperature was held constant with a modified Berendensen thermostat, and pressure was held constant with a Parrinello-Rahman barostat (49). However, the covalent bonds were restrained with the LINCS algorithm (50). The long-range electrostatic energies were computed with Particle Mesh Ewald (PME) method (51) with the cut-off of 1.2 nm. The trajectories from the production run were treated for removing the periodic boundary conditions (PBC) and then used in the analysis. The root mean square deviations (RMSD) from initial equilibrated positions of C-α atoms of each chain of TLRs and respective vaccine chains was investigated to gauge the stability of respective complexes. The fluctuations in the side chain atoms of each residue of TLR chains and vaccine chain was analyzed as root mean square fluctuation (RMSF) in each chain of systems. The compactness of system and consequent stability of system was analyzed in terms of radius of gyration (Rg) for each chain of TLRs and vaccine chain. The hydrogen bonds formed between vaccine chain and the TLR chains were analyzed using appropriate index files of respective chains. Further, the hydrogen bonds formed between vaccine and TLR chains were visually inspected using ChimeraX interface in equilibrated trajectory and trajectory extracted at 25, 50, 75 and 100 ns simulation time. Using the gmx_mdmat program the smallest distance between residues pairs for both the complexes were analyzed and used to obtain the residue wise contact maps (52). Principal component analysis (PCA) (53) was performed to study the major path of motions where the program gmx_covar was employed to obtain a covariance matrix for the C-α atom of each complexes. This covariance matrix was diagonalized using gmx_anaeig program to get the eigenvectors and eigenvalues, where eigenvectors shows the path of motion while the eigenvalues show the mean square fluctuation. The first two principal components (PC1 and PC2) were in Gibb’s free energy landscape (Gibb’s FEL) analysis (54) using gmx sham program. In Gibb’s FEL the deep valleys represent the lowest energy states while the boundaries between the deep valleys show intermediate conformations of the systems. The secondary structural changes were analyzed from definition of protein secondary structure (DSSP) program (55, 56). Further, the extent to which the fluctuations and displacements of a side chains of each TLR chains and vaccine chain are correlated with one another was analyzed from dynamical cross-correlation matrix (DCCM) (56). Poisson Boltzmann surface area continuum solvation (MM-PBSA) calculation (57, 58) were performed to derive the binding free energy estimates between TLR chain and vaccine chain. However, as there are four larger chains in each TLRs and as MM-PBSA calculations are quite extensive, only 10 trajectories which were extracted at each 10 ns from MD simulation run were employed. The protein structures were rendered in ChimeraX (59), PyMOL (60), and VMD (57) and graphs were plotted in XMGRACE (58) interface, while the Gibb’s FEL plots were generated using python based Matplotlib package (61). The DCCM analysis was performed in R statistical program (62) using Bio3D package (63).

### 2.6. Disulfide engineering and *in silico* cloning studies

Vaccine protein disulfide engineering was performed using the Disulfide by Design 2 server (http://cptweb.cpt.wayne.edu/DbD2/) to investigate the conformational stability of folded proteins. Throughout the analysis, the Cα-Cβ-Sγ angle was kept at its default value of 114.6° ± 10, and the χ3 angle was set at -87° or +97°. Residue pairs with energies lower than 2.5 Kcal/mol were chosen and converted to cysteine residues to form disulfide bridges (64). During the in silico cloning study of the vaccine construct, the E. coli strain K12 was selected as the host. However, since the codon usage in humans and E. coli differs, the codon adaptation tool JCAT (http://www.jcat.de) was utilized to adapt the codon usage of the vaccine construct to well-characterized prokaryotic organisms in order to enhance the expression rate (65). When using the JCAT server, it is important to avoid prokaryotic ribosome binding sites, BglII and Apa1 cleavage sites, and Rho independent transcription termination sites. The optimum sequence of the vaccine construct was then inverted, followed by conjugation of the N- and C-terminal BglII and Apa1 restriction sites using the SnapGene Software (66).

## 3. Results

### 3.1. Strain and protein selection with biophysical property analyses

The protein sequences of the target protein (Envelope glycoprotein gp62) in FASTA format were retrieved from the UniProt database. **Table 1** lists the UniProt accession numbers of the selected protein sequences. All the selected proteins were highly antigenic, stable, and have desirable physicochemical properties as listed in **Supplementary Table S1.**

**Table 1:** List of the proteins used in this study with their UniProt accession numbers.

| Strain | Isolate No. | Name of the protein | UniProt accession no. |
| --- | --- | --- | --- |
| HTLV-1 | 01 | Envelope glycoprotein gp62 | P23064 |
|  | 02 | Envelope glycoprotein gp62 | P0C212 |
|  | 03 | Envelope glycoprotein gp62 | Q03816 |
|  | 04 | Envelope glycoprotein gp62 | Q03817 |
|  | 05 | Envelope glycoprotein gp62 | P14075 |
|  | 06 | Envelope glycoprotein gp62 | P03381 |
| HTLV-2 | 01 | Envelope glycoprotein gp62 | P03383 |
| HTLV-3 | 01 | Envelope glycoprotein gp62 | Q0R5Q9 |
|  | 02 | Envelope glycoprotein gp62 | Q09SZ7 |

### 3.2. Epitope mapping for vaccine construction

The vaccine development process included the prediction of T-cell and B-cell epitopes, as well as the assessment of their biophysical characteristics. Subsequently, the epitopes underwent an evaluation to determine their high antigenicity, non-allergenicity, non-toxicity, conservation across selected strains, and dissimilarity to the human proteome. Those that met these criteria were ultimately selected. Additionally, HTL epitopes were further evaluated for their ability to elicit cytokines, and those that could induce at least one cytokine were included in the vaccine. A list of potential CTL, HTL, and LBL epitopes is provided in **Supplementary Table S2**. Eventually, 8 CTL, 7 HTL, and 2 LBL epitopes were chosen based on the stringent criteria mentioned in **Supplementary Table S3** for vaccine construction. Specific linkers and adjuvants were used to conjugate the epitopes, and the vaccine designs were subjected to rigorous testing to confirm their high antigenicity, non-allergenicity, and non-toxicity. **Figures 1A and 1B** illustrate schematic and constructive representations of vaccine construct-1 and 2, respectively, and the conservancy of the epitopes among the selected viral strains is shown in **Figure 2**.

**Figure 1.**
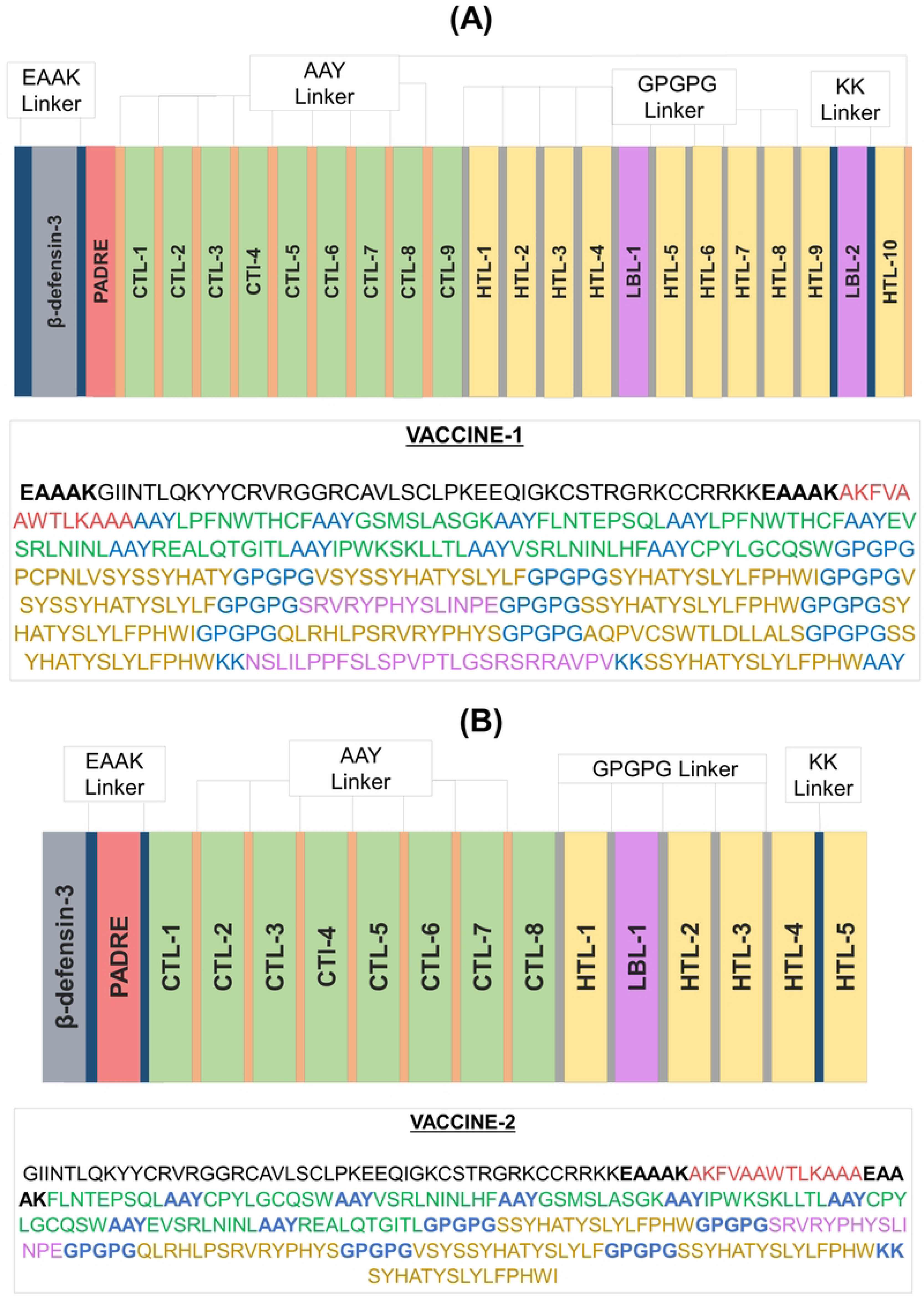
Schematic and constructive representation of (A) vaccine construct 1 and (B) vaccine construct 2

**Figure 2.**
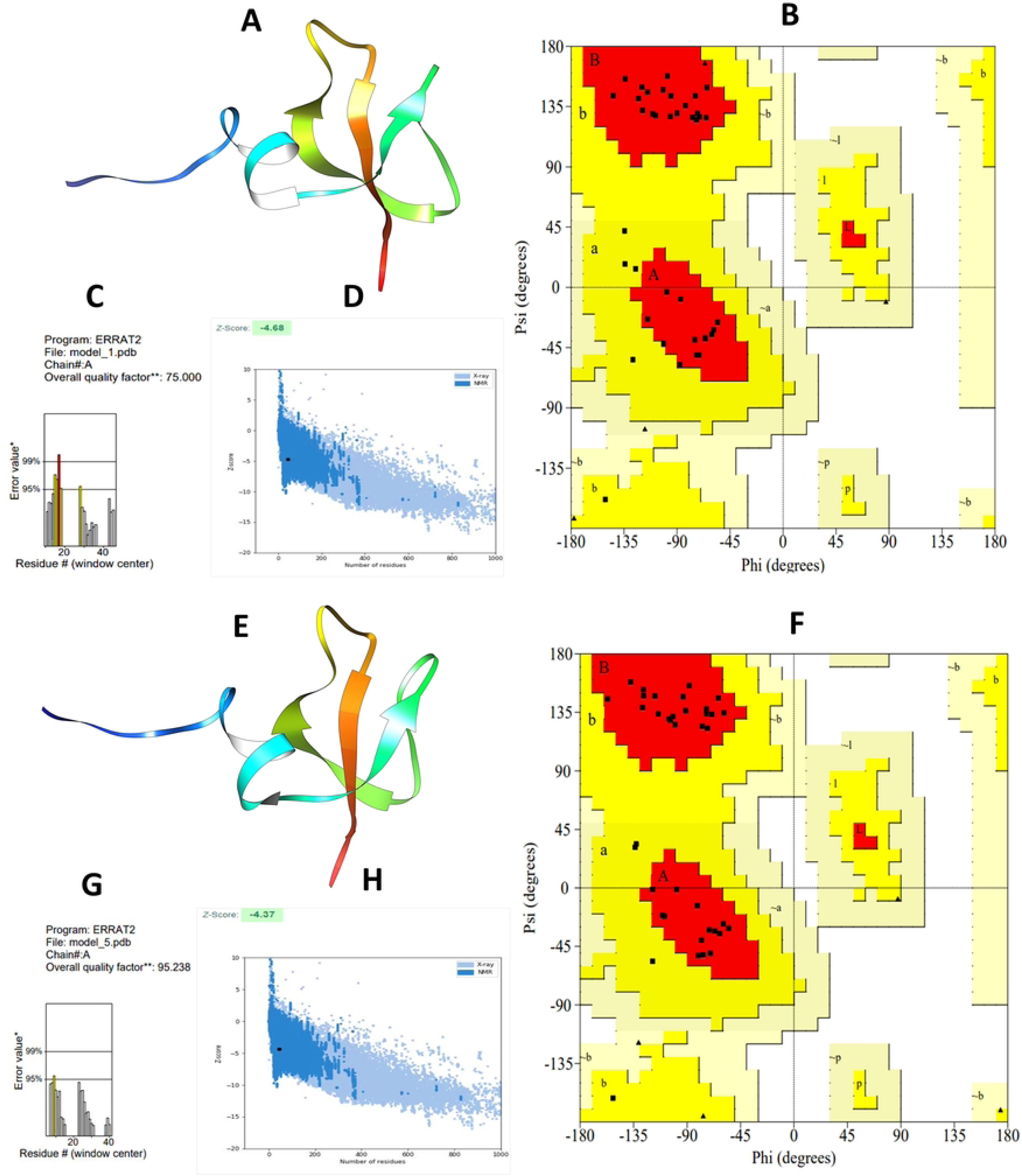
The multiple sequence alignment analysis demonstrating the conservation of the selected epitopes across various strains and isolates of HTLV. The red boxes highlight the presence of these epitopes in multiple HTLV strains, indicating their conserved nature.

### 3.3. Analyses of the biophysical and structural properties of the vaccine

According to biophysical analyses, vaccine construct-1 and 2 displayed favorable qualities such as solubility, stability, and suitability for further examination (as shown in **Supplementary Table S4**). Secondary structure analysis of the vaccine designs indicated that random coil was the most prevalent structure. Afterward, 3D structures of the vaccine constructs were generated, which were subsequently refined and validated. The ERRAT value of vaccine construct-1 and vaccine construct-2 were 75 and 95.238 while the Z score were -4.68 and -4.37 respectively. The Ramachandran plot revealed that most residues for both vaccine designs (84.2% for vaccine construct-1 and 89.5% for vaccine construct-2) were in the favored region. The physicochemical property analysis revealed that vaccine construct-2 has better theoretical Isoelectric point (9.55vs9.63), Aliphatic index (71.60vs73.00), GRAVY (-0.156vs -0.233), and other characteristics in comparison to vaccine construct-1. The overall biophysical and structural characteristics of the vaccines indicated that vaccine construct-2 was more appropriate for further analysis. The 3D model and validation of both vaccine constructs can be found in **Figure 3**, while additional information on the secondary structure of the vaccine constructs is provided in **Supplementary Table S5 and Figure S2**.

**Figure 3.**
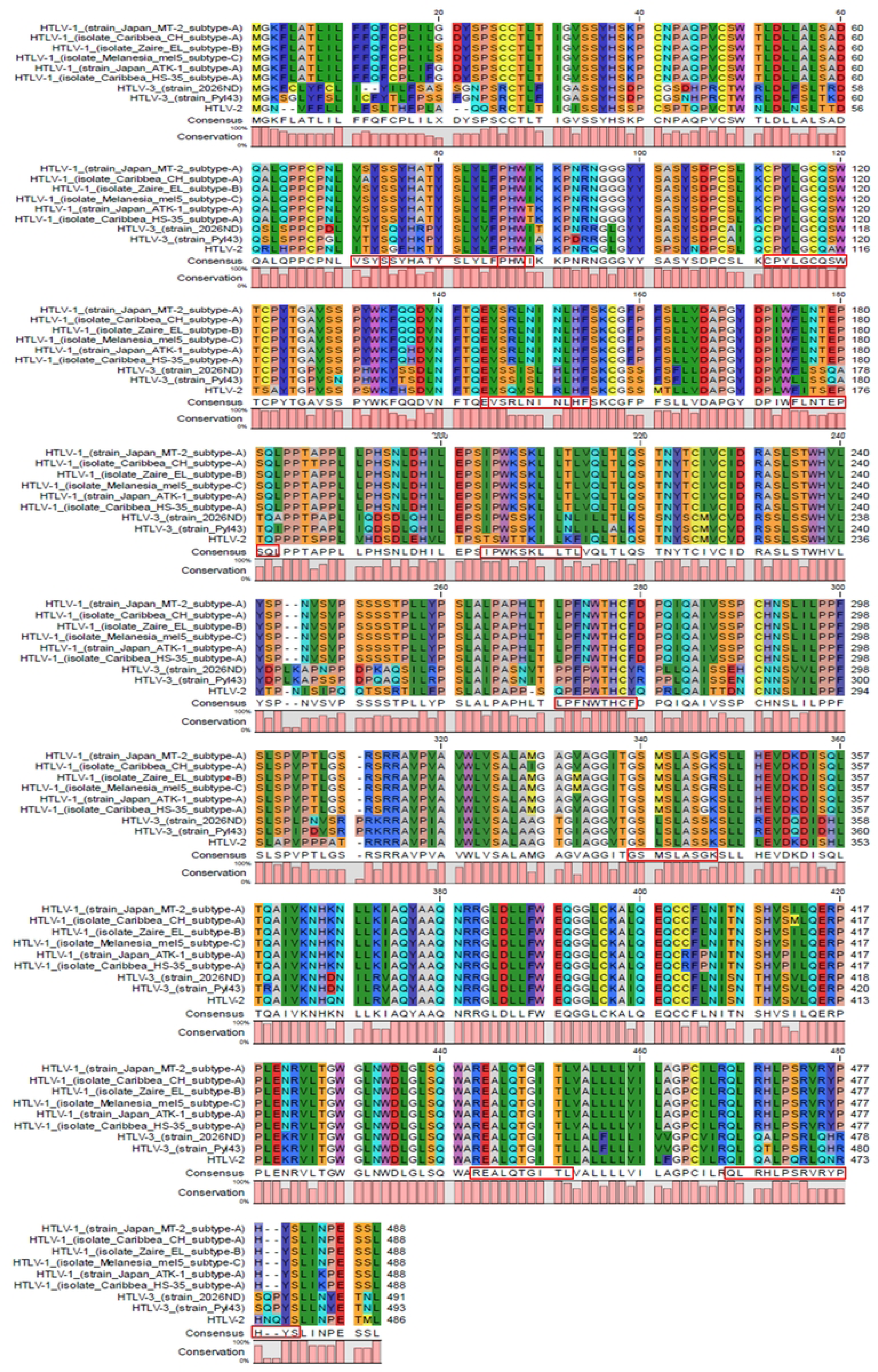
Structure prediction and validation of vaccine construct 1 (A) 3D model (B) Ramachandran layout and (C) The ERRAT quality value (D) Z score graph, and vaccine construct 2 (E) 3D model (F) Ramachandran layout and (G) The ERRAT quality value (H) Z score graph

### 3.4. Molecular docking analyses

To assess the interaction between vaccine epitopes and their corresponding MHC-I alleles, we performed the molecular docking analysis. Since vaccine construct-2 exhibited more promising results in terms of biophysical and structural properties, we specifically focused on conducting the analysis using the CTL epitopes chosen to construct of vaccine 2 and their respective MHC-I alleles. **Supplementary Tables S6 and S7** present the list of epitopes, along with their interacting alleles and docking scores obtained from this analysis. Most of the potential T-cell epitopes exhibited strong binding affinities against both the HLA-A11:01 and HLA-DRB104:01 alleles, as shown in **Figure 4**. Afterward, molecular docking analysis was also conducted to assess the binding affinity of both vaccine constructs with TLR2 and TLR4. The results revealed that vaccine construct-2 had a significantly higher free binding energy and demonstrated a greater docking score with both TLR2 and TLR4. The docking scores obtained from ClusPro and H-dock servers can be found in **Table 2 and Supplementary Table S8**, respectively. Based on the assigned docking score, solubility, and other desired criteria, vaccine construct-2 was selected for additional Molecular Dynamics (MD) simulations using the Gromacs 2020.4 package to further evaluate its interaction and binding affinity with TLR2 and TLR4 (Berendsen, 1995).

**Figure 4.**
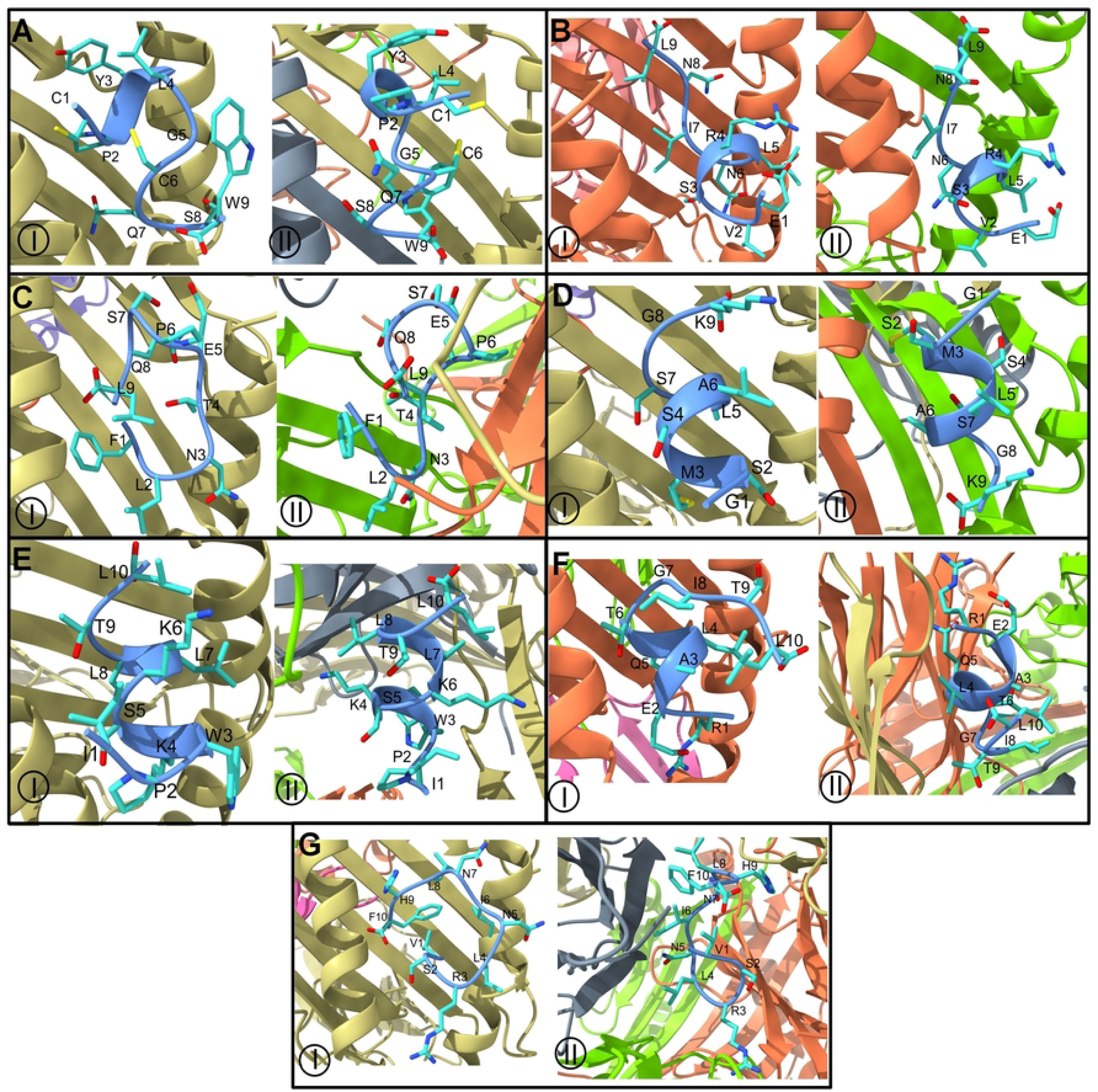
Docked complexes of HTLV-vaccine epitopes. Docked vaccine epitope sequences A) CPYLGCQSW, B) EVSRLNINL, C) FLNTEPSQL, D) GSMSLASGK, E) IPWKSKLLTL, F) REALQTGITL, and G) VSRLNINLHF. (I: HLA-A, II: HLA-DRB; chains of HLA are shown in different colored ribbons, Chain A: light brown, Chain B: light green, chain D: shade of khakhi, chain E: light pink, and vaccine epitopes are shown in light blue ribbon and stick representation)

**Table 2:** Binding affinity between Vaccine molecules and TLRs by ClusPro server.

| TLR's | Vaccine | ClusPro docking energy |
| --- | --- | --- |
| TLR2 | V1 | -763.3 |
|  | V2 | -1018.7 |
| TLR4 | V1 | -869.6 |
|  | V2 | -1054.1 |

### 3.5. Molecular dynamics simulation studies

#### 3.5.1. Root mean square deviation evaluation

The RMSDs in the C-α atom were analyzed independently for each chain of TLRs and vaccine chain. In the TLR2-vaccine complex, chain D has a higher number of fluctuations throughout the simulation period with an average of 0.390 nm (**Figure 5A, Table 3**), however reaching beyond 0.5 nm occasionally. Chain A has a slightly higher RMSD after around 50 ns than chains B and C. The average RMSD for chain A is 0.351 nm, slightly higher than the average RMSD of 0.314 nm for chain C. Chain B has the lowest average RMSD of 0.291 nm with slightly higher deviations during 50-70 ns which is stabilized thereafter. The RMSD for the vaccine chain is slightly higher during the first 10 ns simulation period reaching a maximum of 0.4 nm. Thereafter the RMSD almost remained stable until 90 ns with an average of 0.299 nm and rose to 0.5 nm during last 10 ns simulation period. In the case of the TLR4-vaccine complex, chains C and D almost stabilize after around 15 ns with an average RMSD of 0.288 and 0.260 nm, respectively (**Figure 5B**). Chain A has initially low deviations until about 55 ns which rises to around 0.5 nm till 70 ns and again stabilizes with lower RMSD with an overall average RMSD of 0.265 nm. Chain B has deviations after approximately 10 ns with an average of 0.316 nm. The vaccine chain bound to TLR4 has major deviations until the 60 ns simulation period and stabilized thereafter until the end of the simulation period with an average of 0.344 nm.

**Figure 5.**
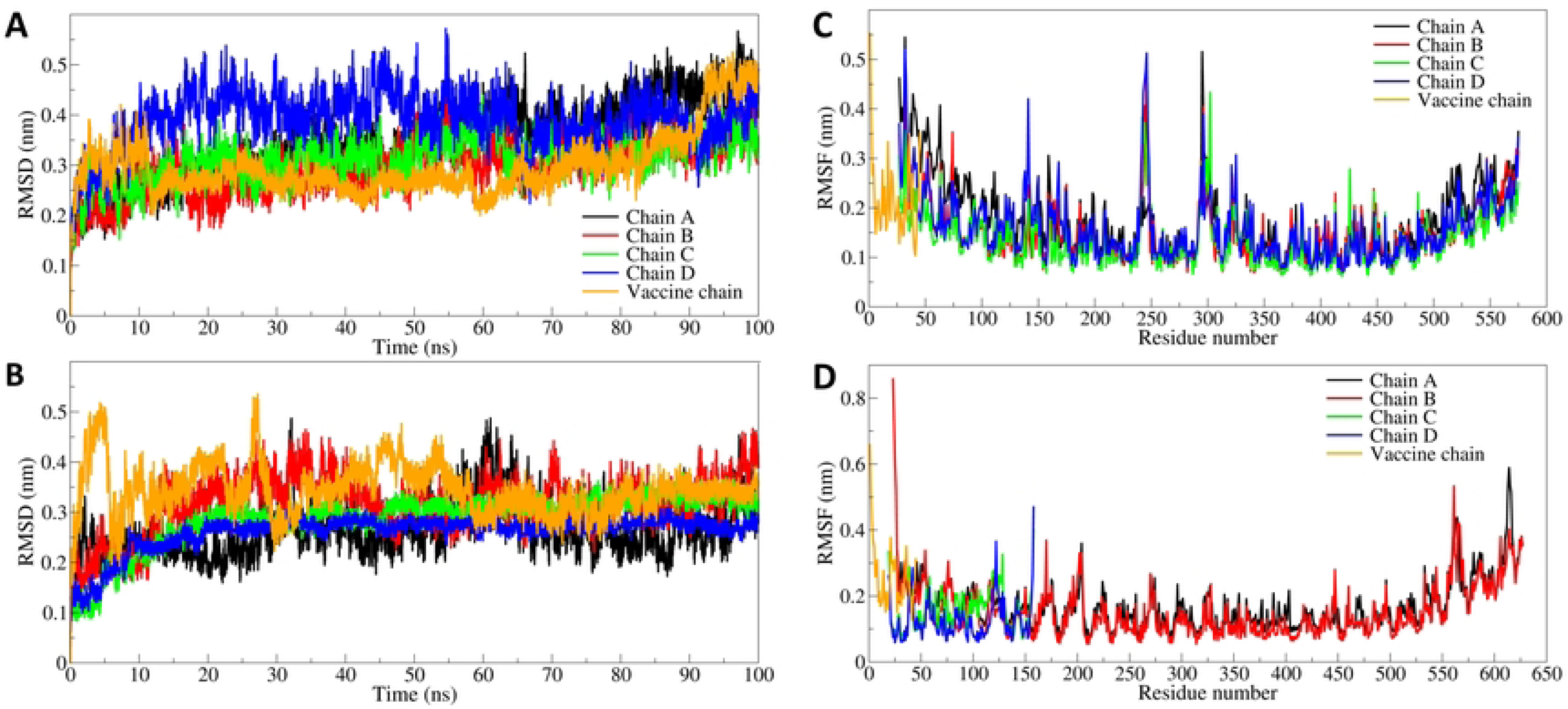
The RMSD analysis. A) RMSD plot for TLR2 chains and bound vaccine chain, and B) RMSD plot for TLR4 chains and bound vaccine chain (color scheme is same as A). The RMSF in side chain atoms of residues. C) RMSF in TLR2 chains and bound vaccine chain, and D) RMSF in TLR4 chains and bound vaccine chain

**Table 3.** Estimates of averages for different MDS analysis parameters

|  | Average (nm) |  |  |
| --- | --- | --- | --- |
| | RMSD in C-α atoms | RMSF | Gyrate |
| TLR2 (Chain A) | 0.351 (0.076) | 0.176 (0.078) | 3.106 (0.045) |
| TLR2 (Chain B) | 0.294 (0.051) | 0.149 (0.060) | 3.064 (0.024) |
| TLR2 (Chain C) | 0.314 (0.044) | 0.137 (0.055) | 3.093 (0.022) |
| TLR2 (Chain D) | 0.390 (0.055) | 0.157 (0.067) | 3.110 (0.031) |
| Vaccine chain bound to TLR2 | 0.299 (0.061) | 0.219 (0.084) | 1.130 (0.023) |
| TLR4 (Chain A) | 0.265 (0.053) | 0.169 (0.073) | 3.191 (0.049) |
| TLR4 (Chain B) | 0.316 (0.057) | 0.154 (0.085) | 3.189 (0.027) |
| TLR4 (Chain C) | 0.288 (0.053) | 0.160 (0.062) | 1.576 (0.010) |
| TLR4 (Chain D) | 0.260 (0.035) | 0.127 (0.061) | 1.544 (0.010) |
| Vaccine chain bound to TLR4 | 0.344 (0.049) | 0.258 (0.099) | 1.131 (0.034) |

Standard deviations in average values are given in parentheses

#### 3.5.2. Root mean square fluctuation evaluation

Root mean square fluctuations (RMSF) were also analyzed separately for each chain of TLRs and respective bound vaccine. The four chains of TLR2 have almost similar lengths and showed nearly equal magnitude fluctuations (**Figure 5C**). All four chains showed a slightly higher magnitude of fluctuations rising beyond 0.5 nm in the residues 225-250 and 275-325. Further, chain A has comparably higher fluctuations in the first 125 residues. The average RMSF for these four chains is in the range of 0.157 to 0.176 nm. In the case of the bound vaccine, almost all the residues have slightly higher RMSF with an average of 0.219 nm.

Chain A and chain B of TLR4 are equal in length and showed almost similar and lower RMSF with averages of 0.169 and 0.154 nm, respectively (**Figure 5D**). Chain C and D of TLR4 are similar and shorter in length, and slightly larger fluctuations are seen in chain C residues with an average RMSF of 0.160 nm. The vaccine chain bound to TLR4 similarly has a larger fluctuation with an average of 0.258 nm.

#### 3.5.3. Radius of gyration evaluation

The total radius of gyration (Rg) of TLR2 showed that almost all TLR2 chains have a stable Rg in the range of 3.064 to 3.110 nm, where chain B showed a lower Rg amongst all TLR2 chains (**Figure 6A**). Chain A and D have higher Rg with averages of 3.106 and 3.110, respectively. The bound vaccine has Rg showed slightly higher Rg during the first 10 ns simulation period which after that stabilizes to a stable Rg with an average of 1.130 nm.

**Figure 6.**
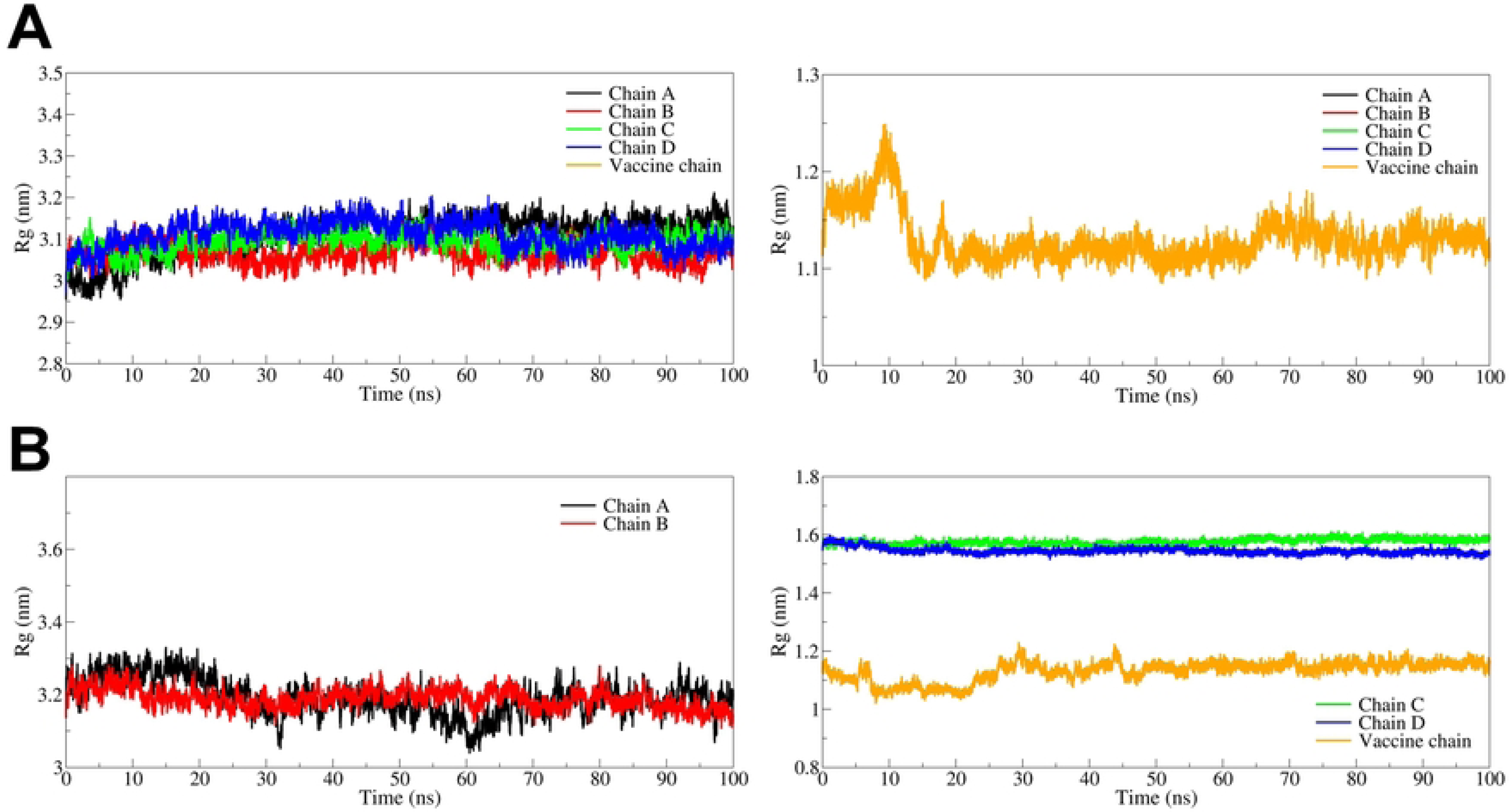
Radius of gyration analysis. A) Rg in TLR2 chains (left hand panel) and vaccine chain (right hand panel), and B) Rg in TLR4 chain A and B (left hand panel) and TLR4 chain C, chain D vaccine chain (right hand panel).

The chain A and B of TLR4 (part of ectodomain) showed average Rg of 3.191 and 3.189 nm, respectively (**Figure 6B**), where the Rg of chain B almost remained stable throughout the simulation. The attached chain C and D (lymphocyte antigenic units) showed slightly lower Rg both of which are stable throughout the simulation with average of 1.576 and 1.544 nm, respectively. The bound vaccine chain showed slight fluctuation until the first 50 ns simulation which stabilized to the lowest Rg with an average of 1.131 nm.

#### 3.5.4. Hydrogen bond analysis

The interchain hydrogen bonds between the vaccine and TLR chains were analyzed. The vaccine chain in the case of TLR2 remained bound at the interface of chains C and D, while in the case of TLR4, it remained at the interface of chains B and D. Thus, the hydrogen bonds formed between vaccine chain and chain C and chain D of TLR2 was first investigated (**Figure 7A**). The vaccine chain was found occupying close contact with TLR2 chain C and an average around 5 hydrogen bonds were found. Initially during the first 25 ns simulation period average of around 5 and a maximum of around 10 hydrogen bonds were found which lowered to average 2 hydrogen bonds till 60 ns and thereafter again around 5 hydrogen bonds re-establish. There is very subtle contact between the vaccine chain and chain D of TLR2 forming only around one hydrogen bond during the first 20 ns simulation period which rises to around an average of 3 hydrogen bonds until 50 ns. However, after 50 ns until the end of the simulation, an average of around 2 hydrogen bonds were seen formed.

**Figure 7.**
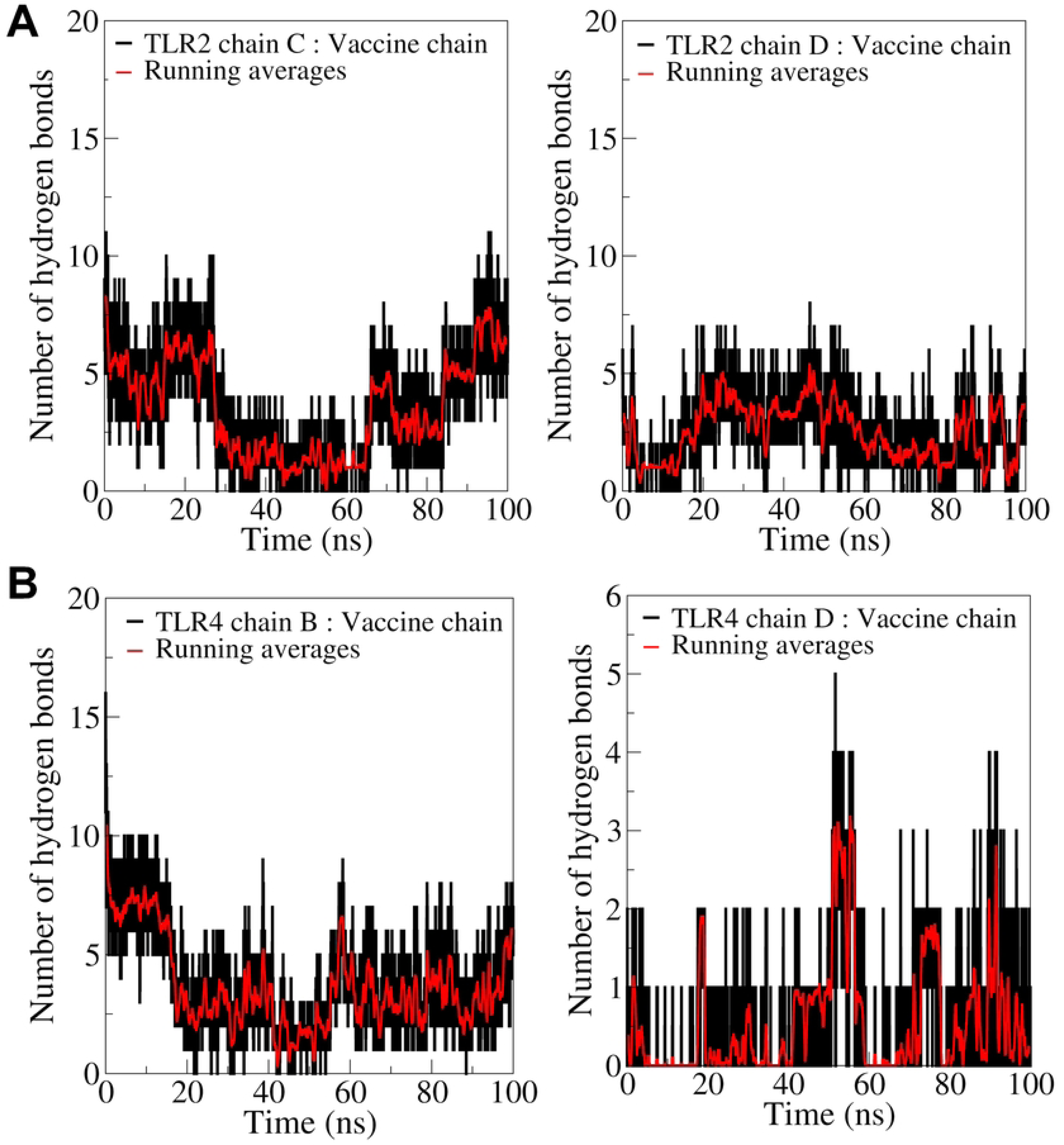
Hydrogen bond analysis plots showing, A) Number of hydrogen bonds formed between TLR2 chain C and vaccine chain (left hand panel) and TLR2 chain D and vaccine chain (right hand panel), and B) Number of hydrogen bonds formed between TLR4 chains B and vaccine chain (left hand panel) and TLR4 chains D and vaccine chain (right hand panel).

Chain B of TLR4 was found in close contact with the vaccine chain and formed around 7 interchain hydrogen bonds during the first 20 ns simulation period reaching a maximum of 10 hydrogen bonds (**Figure 7B**). However, after 20 ns average of only 3 hydrogen bonds were seen formed consistently until the end of the simulation. An average of one interchain hydrogen bond is formed between chain D and the vaccine chain reaching 2 occasionally and reaching a maximum of 5 hydrogen bonds at around 50 ns simulation period.

The interchain hydrogen bonds formed were further visually inspected to investigate important residues of the vaccine chain involved in hydrogen bond formation. The results of this investigation from the equilibrated trajectory and trajectories captured at 25, 50, 75, and 100 ns are briefly presented here. The equilibrated trajectory of TLR2-vaccine complex showed hydrogen bonds between residues Gly1, Ile2, Ile3, Thr5, Leu6, Lys8, Tyr9, Arg12, Val13, Arg14, Thr35, Arg42, and Lys45 of vaccine chain and chain C residues Asp323, Phe322, Asp235, Thr288, Phe325, and Ser329 and chain D residues Asp327, Glu383, Asp384, Asp235, Glu264, Thr236, Leu354, and Gln357 (**Figure 8A**). Except for the hydrogen bond between Lys45 of the vaccine chain and Glu264 of chain C, most of these hydrogen bonds are transient and the trajectory at 25 ns showed that the residues Gly1, Ile3, Asn4, Gln7, Lys32, Thr35 of the vaccine form a hydrogen bond with different residues of chain C *viz.* Asn294, Ser298, Phe325, and chain D residues *viz.* Asn294, Ile304, Ser329, Asn297, and Asp299 (**Figure 8B**). The trajectory at 50 ns showed that the hydrogen bond between the residues Gly1, Asn4, Thr35, and Lys45 from vaccine chain and chain C residue Glu264, and chain D residues Asn294, Ile304, and Asp299 remained stable with no new hydrogen bonds (**Figure 8C**). The trajectories captured at 75 ns showed that the residues Thr10, Lys32, and Lys45 from the vaccine chain form a hydrogen bond with chain C residue Asn294, Ser298, and Asp299 and chain D residue Asp299 (**Figure 8D**). The trajectory at 100 ns showed these hydrogen bonds remained intact and additionally a new hydrogen bonds between residues Gly1, Lys8, Thr9, Arg14, and Arg42 from vaccine chain and chain C residues Asp327, Gly291, Val292, Gly293 and chain D residue Glu336 (**Figure 8E**).

**Figure 8.**
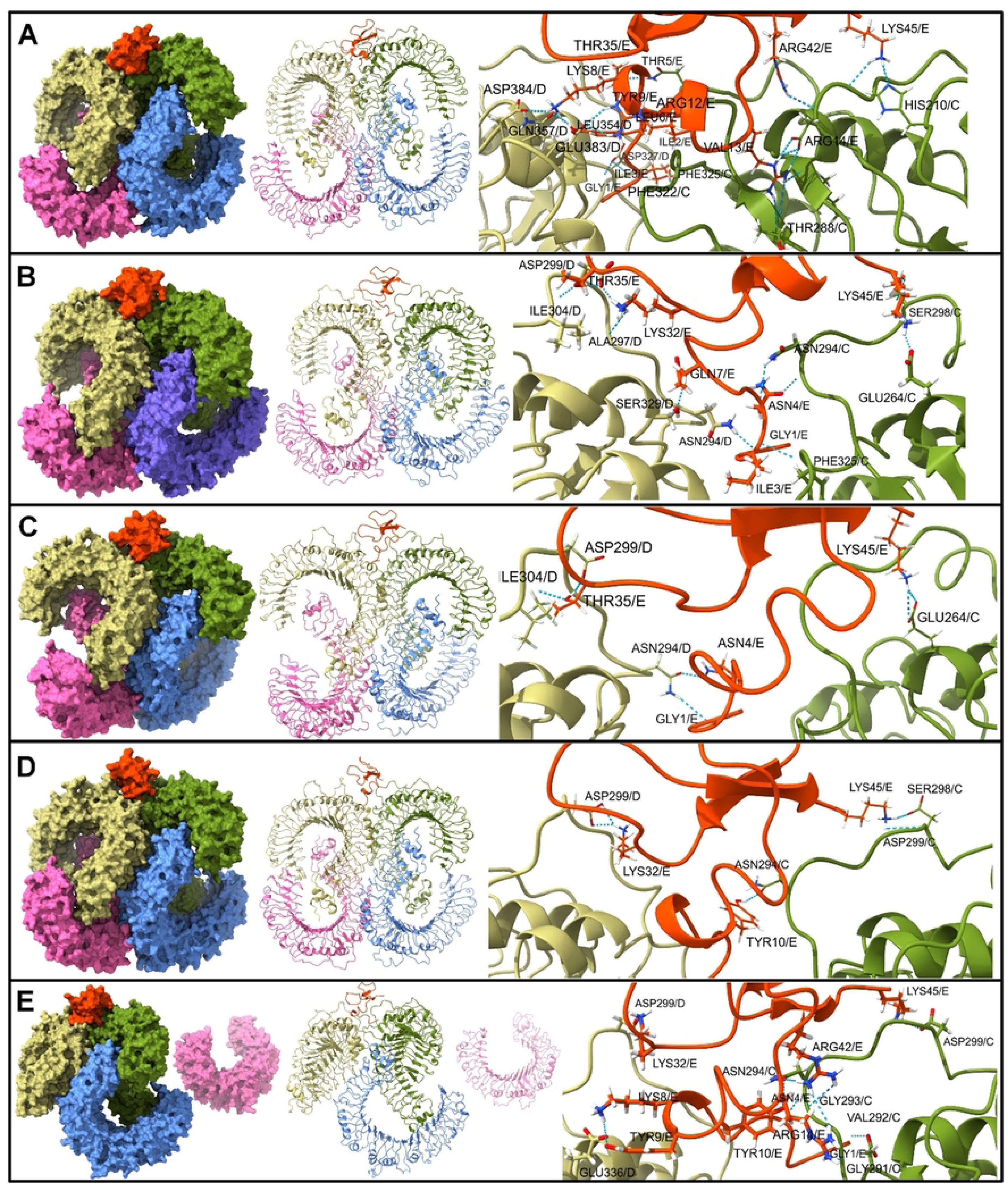
The inter-chain hydrogen bonds between vaccine chain (shown in orange color surface/ribbon/sticks) and two chains of TLR2 *viz.* chain C (shown in olive green surface/ribbon/sticks) and chain D (shown in light brown surface/ribbon/sticks).

In the case of the TLR4-vaccine complex, the initial equilibrated trajectory showed many hydrogen bonds between the vaccine chain and TLR4 B and D chains. The residues Arg12, Arg14, Arg17, Cys18, Ser22, Lys26, Cys33, Ser34, Arg36, Arg38, Arg43, Lys45, and Thr35 formed a hydrogen bond with chain B residues Arg355, Arg382, Glu425, Glu474, Asp502, Asp428, Glu603, Tyr403, Asp379, Glu336, Asp379, Gln523, Asp550, Ser381, Asp405, Gln547 and chain D residue Gln73 (**Figure 9A**). None of these hydrogen bonds are stable, and a new and lone hydrogen bond was formed between the Arg36 residue of the vaccine and Glu287 of chain B (**Figure 9B**). The trajectory at 50 ns showed hydrogen bonds between residues Thr5, Tyr9, and Ser22 of vaccine chain and chain B residues Gln523, Glu474, and chain D residue Ser141 (**Figure 9C**). The trajectory at 75 ns showed the hydrogen bond between Tyr9, Arg17, and Lys45 of the vaccine chain and Glu474, Glu603 of chain B and Ser141 of chain D (**Figure 9D**). The last 100 ns trajectory showed the hydrogen bonds between the residues Thr5, Lys8, Thr9, Arg17, Lys45 and residues Arg606, Gln523, Glu474, and Glu603 of chain B and Ser141 of chain D (**Figure 9E**).

**Figure 9:**
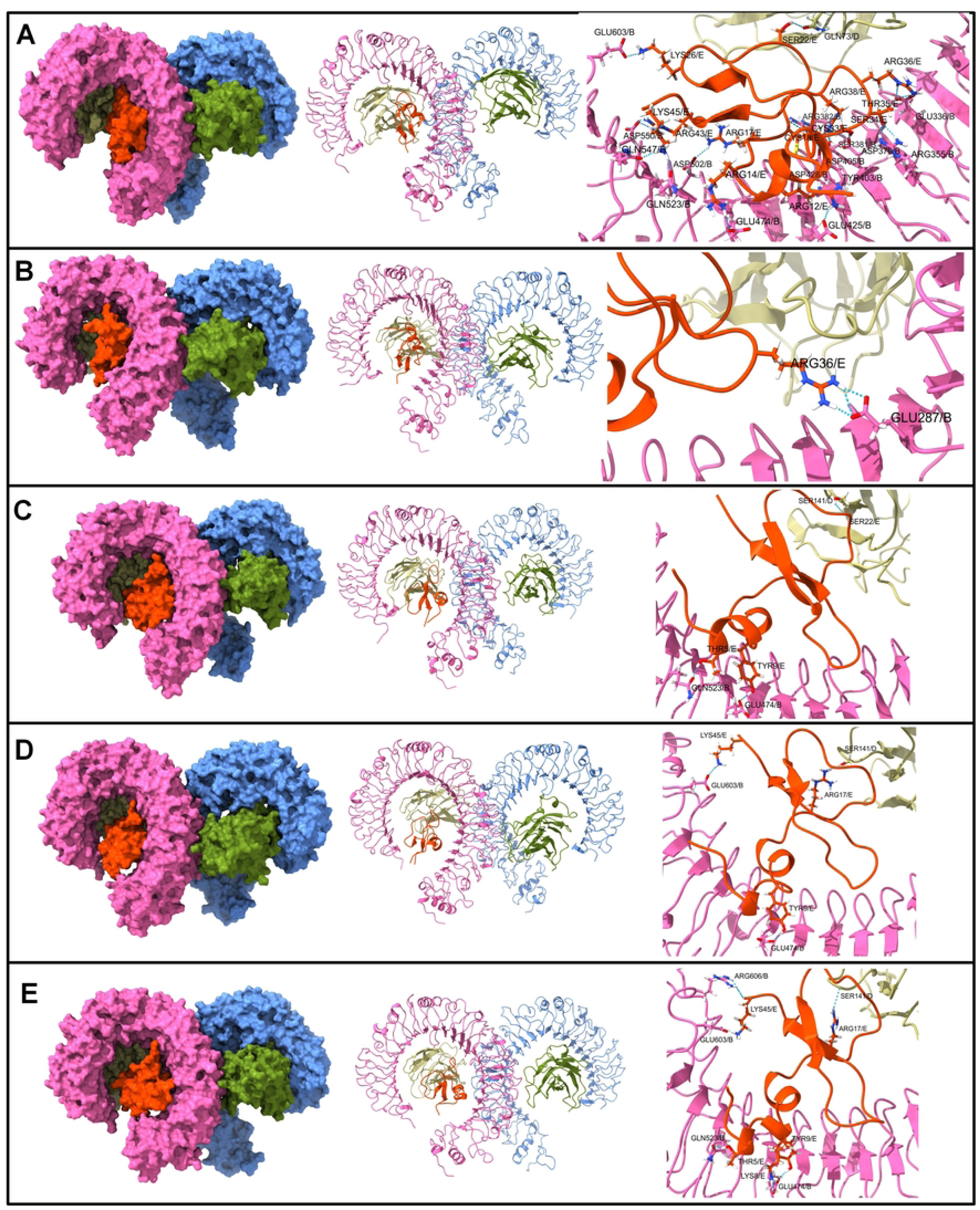
The inter-chain hydrogen bonds between vaccine chain (shown in orange color surface/ribbon/sticks) and two chains of TLR4 *viz.* chain B (shown in pink surface/ribbon/sticks) and chain D (shown in light brown surface/ribbon/sticks).

#### 3.5.5. Contact map analysis

In the TLR2 vaccine complex, the vaccine is bound at the interface of chains C and D. While in the TLR4 vaccine complex, the vaccine is bound at the interface of chains B and D. The extent of residue-residue contacts between these TLR chains and the vaccine chain was analyzed through contact map analysis (**Figure 10**).

**Figure 10.**
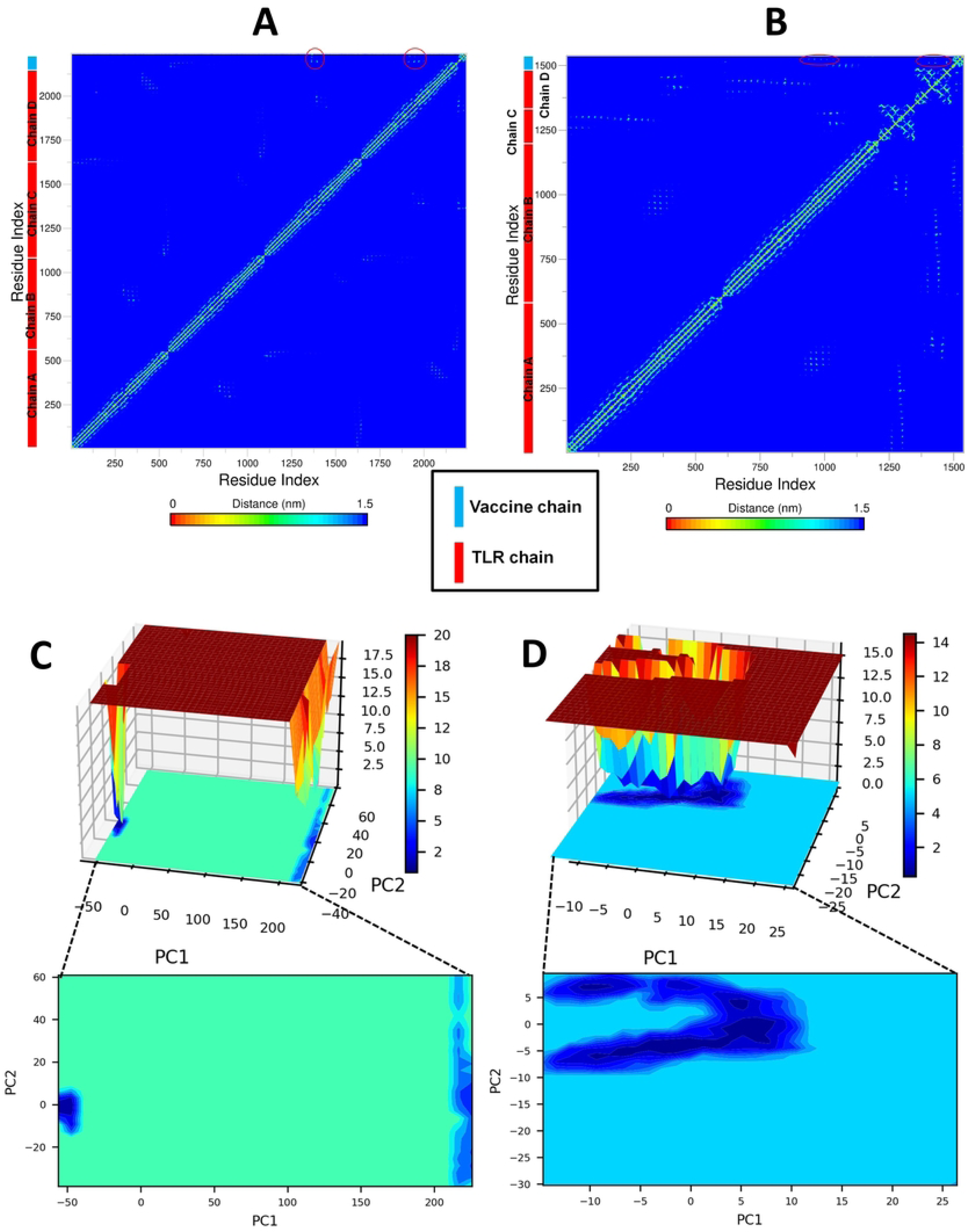
Contact maps constructed for the mean smallest distances between network of residues of TLR2 and vaccine. A) TLR2-vaccine complex, and B) TLR4-vaccine complex (The residue-residue contacts between respective TLR chains and vaccine chain are marked in red circles). Gibb’s free energy landscape. C) TLR2-vaccine complex, and D) TLR4-vaccine complex

The contact maps results show that the TLR2 chain C has slightly a greater number of residue-residue contacts with the vaccine chain compared to residue-residue contacts between chain D and the vaccine chain. More residue-residue contacts were found between TLR4 chain B, and the vaccine chain compared to residue-residue contacts between TLR4 chain D and the vaccine chain (**Figure 10A-B**).

#### 3.5.6. Principal component analysis and Gibb’s free energy analysis

Principal component analysis (PCA) and corresponding Gibb’s free energy analysis were performed. Comparably, a large energy basin with the lowest energy was found with the TLR4-vaccine complex (**Figure 10**). Among the two energy basins in the TLR2-vaccine complex, the one occupying the energy range of around -50 kJ mol^-1^ on PC1 and 5 to -15 kJ mol^-1^ on PC2 is the lowest energy basin, while the other one with the higher energy range is beyond 200 kJ mol^-1^ on PC1 and spread across 60 to -30 kJ mol^-1^ on PC2 (**Figure 10C**). The TLR4-vaccine complex has a unique energy basin spread across 10 to -20 kJ mol^-1^ on PC1 and 10 to -10 kJ mol^-1^ on PC2, suggesting a large stable lowest energy conformation (**Figure 10D**).

#### 3.5.7. Dynamic cross-correlation (DCC) analysis

Through DCC analysis, the time-correlated information of inter-chain and intra-chain residue to residue contacts and motions was analyzed. In DCC plots, the color gradient ranges from blue (negative correlation, less likely) to red (positive correlation, more evident) corresponding to the correlation coefficients -1 and +1, respectively, and the lighter shades indicate weaker correlations, where the white color indicates no correlation. The DCC correlation of the vaccine chain with each chain of TLRs was analyzed (**Figure 11A-B**). The results for the TLR2-vaccine complex indicate that the residues from chains B, C, and D are positively correlated with the vaccine chain with a stronger correlation with chains B and C. Similarly, in the TLR4-vaccine complex, chains B, C, and D have moderate to strong positive correlation with vaccine chain.

**Figure 11.**
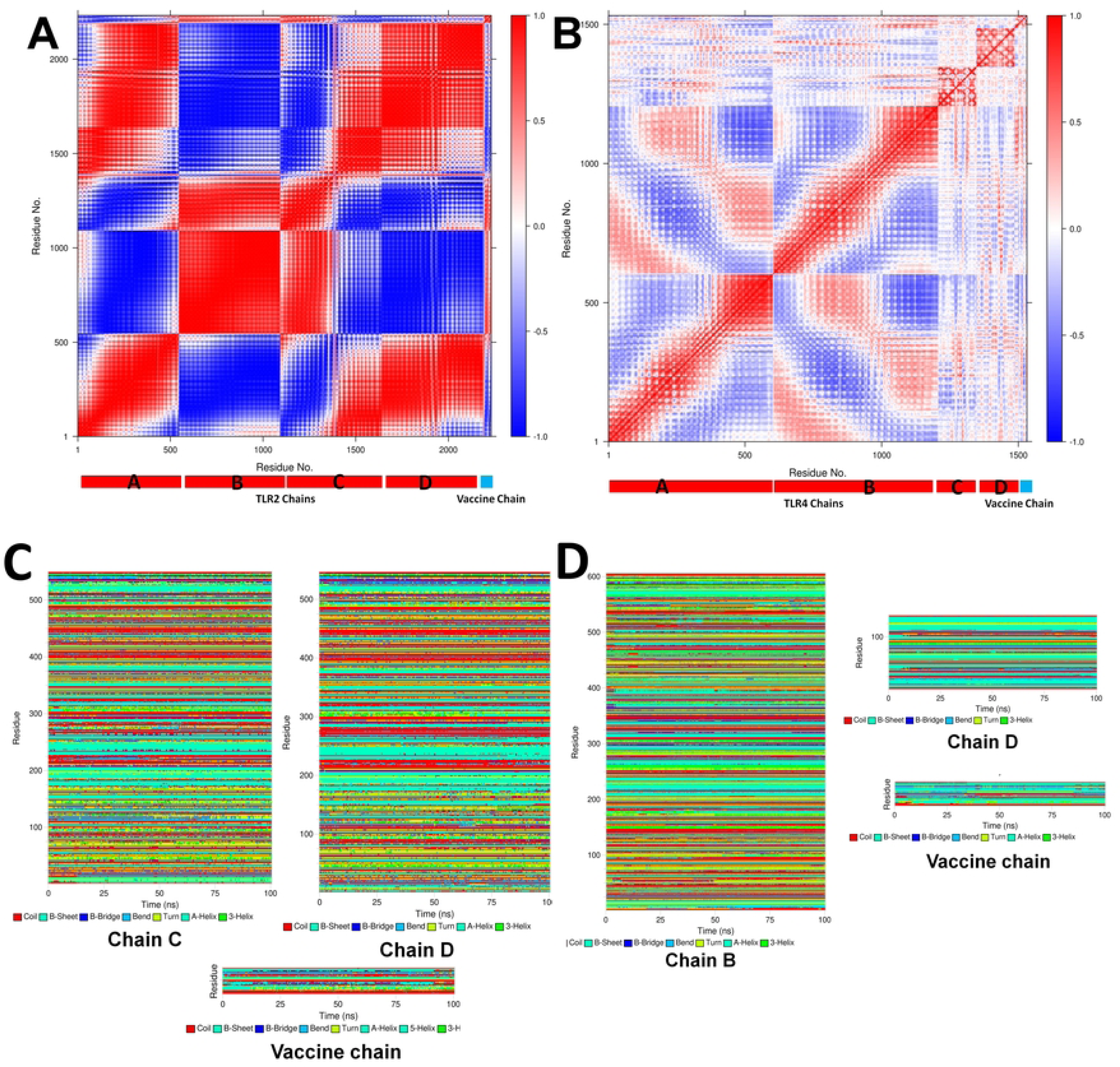
DCC analysis for vaccine chain complex with A) TLR2, and B) TLR4. (The plot of DCCM shows residue-wise correlation in each complex. The separation of TLR chain and vaccine chain is shown at the bottom of each plot). DSSP plots for individual TLR chains and corresponding vaccine chain. C) TLR2-vaccine complex and D) TLR4-vaccine complex

#### 3.5.8. DSSP analysis

The secondary structural changes which occur during MD simulation were analyzed from DSSP analysis. Especially, the TLR chains which are in close contact with vaccine chains were analyzed. The results showed that the TLR2 chain C and chain D are relatively stable for secondary structural changes (**Figure 11C-D**). However, the vaccine chain showed major structural changes in the loop regions. Similarly, chains B and D of TLR4 showed quite stable secondary structure, while the vaccine chain bound to TLR4 showed numerous secondary structural changes in the loop regions.

#### 3.5.9. MM-PBSA calculation

The MM-PBSA calculations on 10 trajectories isolated at each 10 ns for each complex were performed. In MM-PBSA calculations, the interaction energies *viz.* van der Waal energy, electrostatic energy, polar solvation energy, SASA energy, and binding energy (ΔG_binding_) between TLR chains and vaccine chain were estimated (**Table 4**). The results show that the TLR4-vaccine complex has more favorable binding energy compared to the TLR2-vaccine complex. The lower electrostatic, van der Waal, and polar salvation energy for the TLR4-vaccine complex might be responsible for the lower and favorable binding free energy.

**Table 4.**
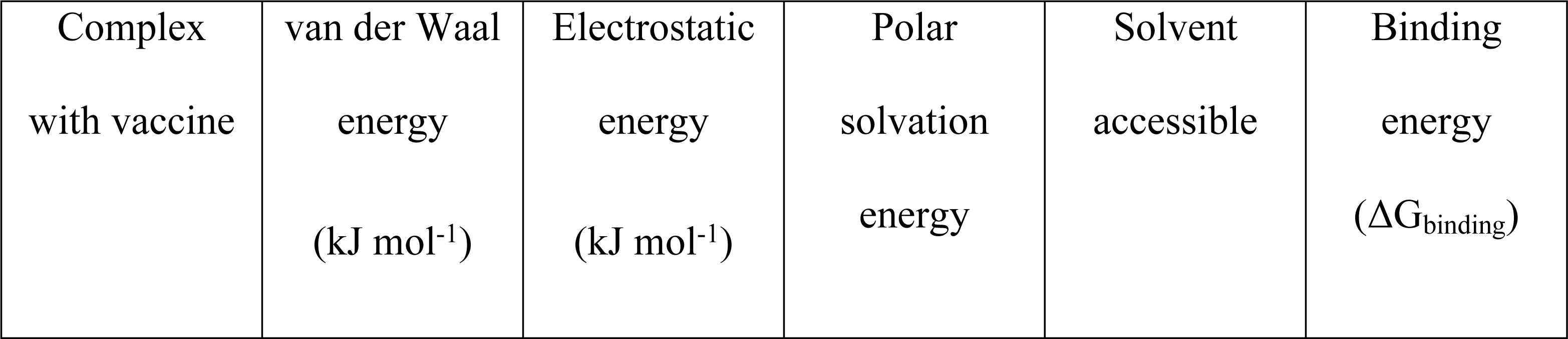
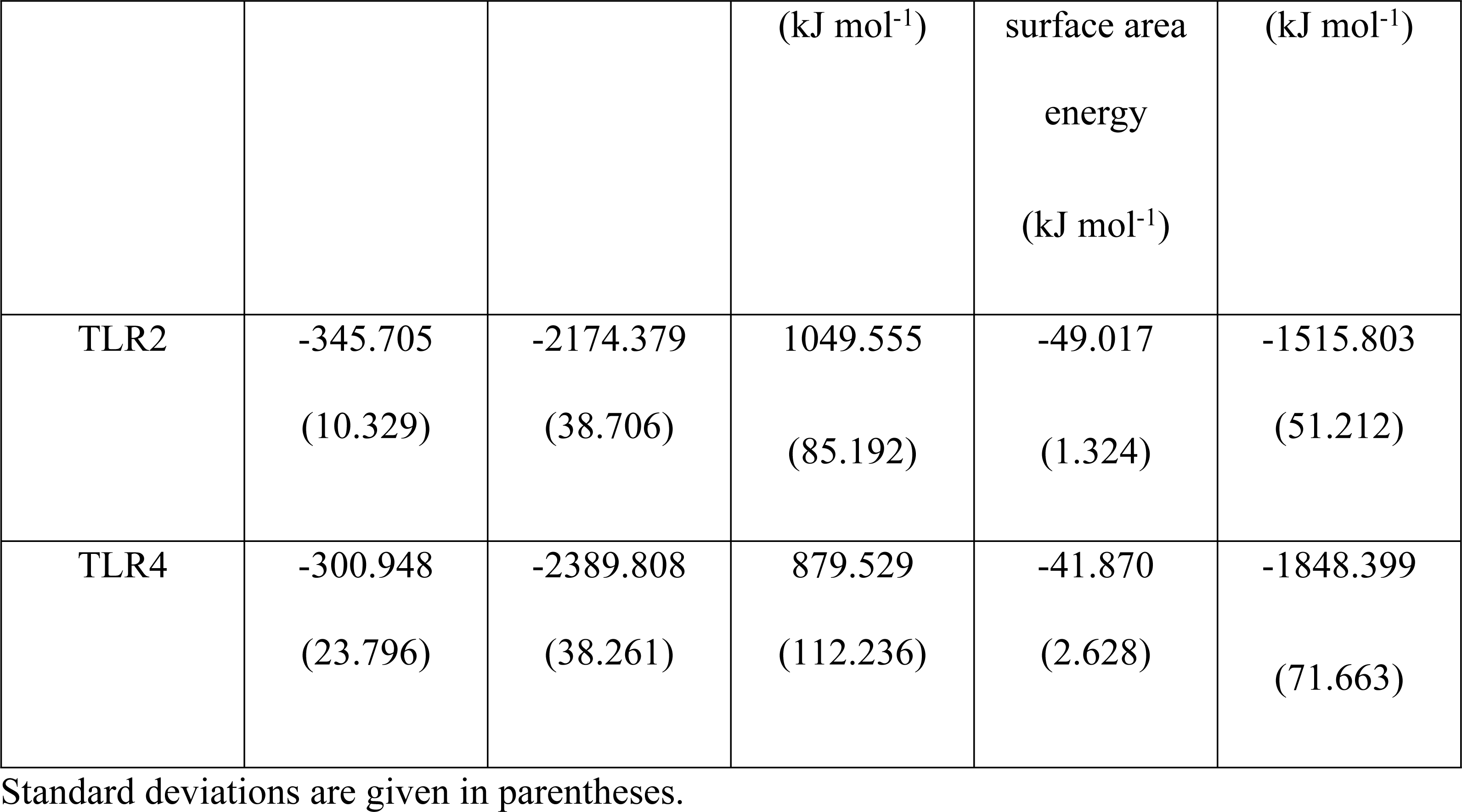
Results of MM-PBSA calculations

### 3.6. Disulfide engineering and *in silico* cloning studies

By using the DbD2 server, a total of 7 pairs of amino acid residues for vaccine construct-2 have been discovered as having the ability to structure disulfide bonds. Three pairs of amino acid residues (Cys 11 – Gly 15, Cys 11 – Cys 40, and Cys 18 – Cys 33) were carefully chosen due to their compatibility with standard disulfide bond formation conditions, with energy levels lower than 2.5 Kcal/mol (**Supplementary Figure S3**). In the codon adaptation study of the vaccine, the codon adaptation index (CAI) revealed that the adapted codons displayed a higher proportion of the most abundant codons. Notably, the optimized codons exhibited a significant GC content of 54.6428 and a CAI of 0.9690. To ensure the safety of the cloning process, the absence of restriction sites for BsgI and AvaI was confirmed. Subsequently, the optimized codons, along with the BsgI and AvaI restriction sites, were inserted into the pET28a (+) vector. This resulted in the generation of a 3515 base pair clone, which included the desired 387 bp sequence, with the remaining portion belonging to the vector. The desired area between the pET28a (+) vector sequence was indicated in red, as depicted in **Supplementary Figure S4**.

## 4. Discussion

Given the devastating impact of the HTLV viruses in causing a diverse group of inflammatory, immunosuppressive disorders and cancer, there is an urgent demand for effective preventive and treatment approaches to mitigate severe illness and mortality. To address this pressing need, our study employed a reverse vaccinology approach to design a multi-epitope-based vaccine that specifically targets the three highly virulent subtypes of HTLVs. Furthermore, we conducted a detailed analysis into the molecular interactions between the vaccine and TLRs, revealing valuable insights that could guide future research on adopting effective strategies and formulating countermeasures to prevent and manage HTLV-related diseases or outbreaks.

The vaccine was designed by targeting the envelope glycoprotein gp62, an outer membrane protein having a greater number of epitope binding site that plays a significant role in the infectious process and immune response of HTLV (67, 68). The selected proteins were predicted to be antigenic and having desirable biophysical properties that are expected in vaccine development. Epitope mapping was conducted to identify both T cell and B cell epitopes as both are crucial for stimulating the host’s immune system in response to viral infection (69, 70). IEDB algorithms were used to predict the epitopes, and all of the chosen epitopes showed low binding energies assuring a higher affinity for their targets. The final epitopes selected for vaccine development were highly antigenic, non-allergenic, and non-toxic, indicating their potential to trigger a sound and safe immune response against the viral infection. The selected T cell and B cell epitopes were connected using appropriate linkers, while also incorporating PADRE and hBD adjuvants to enhance their immunogenicity within the human body. The hBD adjuvant plays a role in promoting innate and adaptive host defense through its ability to chemo-attract immature dendritic cells, naive memory T cells, and monocytes to the site of infection. Moreover, the use of adjuvant and linkers improves the vaccine’s antigenicity, immunogenicity, and lifespan profile while stabilizing the vaccine construct (71).

Two vaccines were developed using the identified epitopes, and both were found to be highly antigenic, non-allergenic, and non-toxic, confirming their safety and potential to trigger a sufficient immune response. Vaccine construct 2 was found to be soluble while vaccine construct 1 was insoluble. Solubility is a vital feature of vaccine since an insoluble vaccine possesses the risk of being ineffective while producing an abundance of insoluble tangled peptide mass inside the body (72). The biophysical property analysis of the vaccine revealed its competence. Both the vaccine construct was found to be stable having an instability index lower than 40. The aliphatic index value, which assesses thermostability, were 71.60 and 73.00 for vaccine construct 1 and vaccine construct 2, respectively, suggesting that the vaccines were stable at normal human body temperature. The negative GRAVY value of the vaccine constructs refers to their hydrophilicity which indicates their higher solubility in water. Furthermore, disulfide engineering in vaccine construct 2 resulted in the generation of three pairs of amino acids that can be mutated into cysteine residues, thereby enhancing the stability and efficacy of the vaccine (73).

Both the vaccine construct had a prominent random coil secondary structure. According to tertiary structure prediction and validation, vaccine construct 2 had a superior structure profile than vaccine construct 1, with a higher ERRAT score (95.238 vs. 75) and more structure in the favored region (89.5% vs. 84.2%). Both vaccine construct showed no structure in the disallowed region of the ramachandran plot assuring higher quality of the vaccine models. Taken together, structure prediction conferred that the both vaccine constructs have appropriate structure and should be sufficiently stable.

During the molecular docking analysis, the CTL epitopes of vaccine construct 2 were initially docked against HLA-A11:01 and HLA-DRB104:01 to evaluate their binding affinity with representative HLA alleles. HLA-A11:01 is a well-known human leukocyte antigen, while HLA-DRB104:01 is commonly found in patients with severe extra-articular rheumatoid arthritis associated with HTLV (74, 75). Both of these alleles play a crucial role in presenting viral antigens to T cells and triggering an effective immune response (76). The epitopes showed high binding affinity with both HLA alleles. Subsequently, molecular docking was performed again to assess the interaction between both vaccine constructs and TLR2 and TLR4 using the ClusPro server and H-dock server. TLR2 and TLR4 are expressed on the cell surface as well as intracellularly in dendritic cells (DCs), endothelial cells, and epithelial cells (ECs), making them suitable targets for vaccine constructs (77). Vaccine construct 2 exhibited the lowest docking scores with both TLR2 and TLR4 in the ClusPro server (-1018.7 and -1054.1, respectively) and H-dock server (-281.88 and -279.46, respectively). Based on the performances of two vaccine constructs in the aforementioned experiments i.e., antigenicity, solubility, physicochemical properties, and binding affinity, vaccine construct 2 was selected for further molecular dynamic simulation studies.

TLR2 has four equal in length chains, chain A, B, C, and D. While, TLR4 has two chain which belong to TLR ectodomain, chain A and chain B and two antigen units, chain C and chain D. The highest magnitude of RMSD was observed in chain D of TLR2, while RMSD in other three chains is almost stable. On the other hand, the RMSD in corresponding chain A and chain B is slightly deviating throughout the simulation. The RMSD in antigen units of TLR4 representing chain C and chain D is quite stable. Further, the RMSD in vaccine chain bound to TLR2 is lower compared to the vaccine chain bound to TLR4. TLR2-vaccine complex seems more stable compared to TLR4-vaccine complex. The RMSF in all the chains of TLR2 are almost analogous and relatively lower except for residues in the range 200-325 suggesting uniform conformational changes in these chains. In the case of TLR4 two ectodomain chains chain A and B has analogous and minimal fluctuations in almost all the residues suggesting very stable conformation of these chains. The antigen units, chain C and D has few fluctuations and especially chain D has mode fluctuations which is in close contact with vaccine chain. In both the complexes the vaccine chain has shown marginal fluctuations in the non-terminal residues suggesting comparable stability of the respective complexes. It is evident that all the four chains of TLR2 have almost similar total Rg suggesting similar compactness of each chain. While two TLR chains, chain A and chain B, from TLR4 showed almost similar compactness of the chains which suggests their analogous compactness and rigid structural features. The antigen units, chain C and D, bound to TLR4 being short proteins also showed very compact nature as they have significantly lower total Rg than TLR4 A and B chains. After around 25 ns the vaccine chains bound to TLR2 and TLR4 suggested very compact nature from the stable and lowest total Rg. The hydrogen bond analysis suggested the vaccine chain in both the complexes forms maximum and more consistent hydrogen bonds with chain C and chain B in the case of TLR2 and TLR4 respectively. However, the hydrogen bonds between chain D and vaccine in TLR2 and TLR4 complex significantly differed. Here, the vaccine chain could form more number and consistent hydrogen bonds with chain D in TLR2 complex. These results suggest that the vaccine chain interact and forms a good number of key hydrogen bonds with chain C and D of TLR2 compared to a vaccine forming hydrogen bonds with chain B and D of TLR4. More the number of non-bonded interactions such as hydrogen bonds better is the affinity of vaccine and overall stability of corresponding complex. On these grounds

TLR2-vaccine complex seems to have better stability and the vaccine construct in this case is having more favorable affinity for the TLR2. The contact analysis also corroborates the results of hydrogen bond analysis where the TLR2-vaccine complex has a greater number of residue-residue contacts between chain C and chain D of TLR2 and vaccine chain. On the other hand, fewer residues of TLR4 chain B and D could establish such contacts with vaccine residues. The existence of low energy conformations from the simulation was analyzed from Gibb’s free energy landscapes. In TLR2-vaccine complex very small low energy basin with energy around -60 to -50 kJ mol-1 nm-1 on PC1 and around -10 to 5 kJ mol-1 nm-1 on PC2 suggests existence of fewer low energy conformations compared to TLR4-vaccine complex. In TLR4-vaccine complex a major low energy basis, where many lowest energy conformations exist, occur at energy range 10 to -20 kJ mol-1 nm-1 on PC1 and 0 to -10 kJ mol-1 nm-1 on PC2. The Gibb’s free energy analysis points out that despite lower number of hydrogen bonds and key contacts the TLR4-vaccine complex has numerous low energy states suggesting more stable state. The DCC analysis revealed that vaccine bound to TLR2 has positively correlated residues from chain B, chain C and chain D and very few from chain A. On the other hand, TLR8-vaccine complex showing moderately positive but many such positively correlated residues from all the chains. There are very few negatively correlated residues in TLR4-vaccine chain which further confirms its better stability. The DSSP analysis points out that the TLR chains viz. chain A and chain D in TLR2 and chain B in TLR4 are quite stable for any secondary structural changes. However, the chain D, the antigen unit in TLR4, undergoes few major secondary structural changes suggesting its structural flexibility. The vaccine chains in both the complexes were seen undergoing secondary structural changes mostly in the loops or turns which might have resulted in better and compact structure of vaccine construct. The MM-PBSA analysis suggested that the vaccine construct bound to TLR4 has more favorable binding affinity compared to the vaccine construct bound to TLR4. The major difference found in the energetic is in electrostatic energy in TLR4 which is significantly lower than TLR2.

Overall, molecular dynamics simulations showed enhanced stability and favorable affinity in the TLR2-vaccine complex, supported by contact analysis and secondary structure stability, while MM-PBSA analysis favored the TLR4-vaccine complex due to lower electrostatic energy.

## 5. Conclusion

The research on the multi-epitope-based vaccine targeting highly virulent subtypes of HTLV has significant future implications. The findings contribute to the development of an effective preventive and therapeutic approach for HTLV-associated disorders. By utilizing a reverse vaccinology strategy, the study demonstrates the design and characterization of vaccine constructs with high antigenicity, safety, and stability. The incorporation of adjuvants and linkers enhances the immunogenicity of the vaccine. Molecular docking studies and molecular dynamic simulations provide insights into the strong binding affinity and stability of the vaccine with key immune receptors, such as HLA alleles, TLR2, and TLR4. These findings lay the foundation for potential strategies to prevent and manage HTLV-related diseases and outbreaks, offering promising prospects for improved healthcare outcomes in the future.

## 7. Declarations

### Data availability statement

All the data generated during the experiment are provided in the manuscript/supplementary material.

### Ethics approval statement

Not Applicable

### Patient consent statement

Not Applicable

### Permission to reproduce material from other sources

Not Applicable

### Clinical trial registration

Not Applicable

### Consent for participation/publication

Not Applicable

### Authors’ contributions

M.A.U. and A.T.M. conceptualized the study. A.T.M., N.A.R., R.B.P., T.B.R., T.Z. and M.A.U. wrote the manuscript draft. R.B.P., A.T.M., N.A.R., M.N.S. and T.B.R. conducted the experiments. R.B.P., A.T.M., and N.A.R. generated the figures. All the authors reviewed the manuscript. M.A.K., N.A., and A.M.S. supervised the study.

## Acknowledgements

We express our sincere gratitude to all those who have supported us throughout our research endeavor. We dedicate this study to Alhajj Muhammad Abu Taher and Syeda Jasmin Akter for their unwavering support, affection, and motivation. We also pay tribute to the late Syed Muhammad Nazmul Hoque and the late Muhammad Abul Bashar, who have been a profound source of inspiration for us. Our deepest gratitude goes to the Laboratory of Clinical Genetics, Genomics, and Industrial Biotechnology at the Department of Genetic Engineering and Biotechnology, University of Chittagong, Chattogram, Bangladesh. We extend our thanks to Mrs. Ayesha Siddique, Ms. Nafisa Nawal, and Mr. Talha Zubair for their support and encouragement throughout our research journey. We are also grateful to Barrister Mohibul Hasan Chowdhury, Deputy Minister of Education of Bangladesh, for his continuous inspiration and encouragement, urging us to strive for excellence in our endeavors. Finally, we dedicate this study to all aspiring student researchers in Bangladesh, whose passion for science and research will undoubtedly lead to significant contributions in the field.

